# Single-cell Transcriptomic Landscape of Nucleated Cells in Umbilical Cord Blood

**DOI:** 10.1101/346106

**Authors:** Yi Zhao, Xiao Li, Jingwan Wang, Ziyun Wan, Kai Gao, Gang Yi, Xie Wang, Jinghua Wu, Bingbing Fan, Wei Zhang, Fang Chen, Huanming Yang, Jian Wang, Xun Xu, Bin Li, Shiping Liu, Weihua Zhao, Yong Flou, Xiao Liu

## Abstract

Umbilical cord blood (UCB) transplant is a therapeutic option for both pediatric and adult patients with a variety of hematologic diseases such as several types of blood cancers, myeloproliferative disorders, genetic diseases, and metabolic disorders. However, the level of cellular heterogeneity and diversity of nucleated cells in the UCB has not yet been assessed in an unbiased and systemic fashion. In the current study, nucleated cells from UCB were subjected to single-cell RNA sequencing, a technology enabled simultaneous profiling of the gene expression signatures of thousands of cells, generating rich resources for further functional studies. Here, we report the transcriptomic maps of 19,052 UCB cells, covering 11 major cell types. Many of these cell types are comprised of distinct subpopulations, including distinct signatures in NK and NKT cell types in the UCB. Pseudotime ordering of nucleated red blood cells (NRBC) identifies wave-like activation and suppression of transcription regulators, leading to a polarized cellular state, which may reflect the NRBC maturation. Progenitor cells in the UBC also consist two subpopulations with divergent transcription programs activated, leading to specific cell-fate commitment. Collectively, we provide this comprehensive single-cell transcriptomic landscape and show that it can uncover previously unrecognized cell types, pathways and gene expression regulations that may contribute to the efficacy and outcome of UCB transplant, broadening the scope of research and clinical innovations.

## INTRODUCTION

Human umbilical cord blood (UCB) is an excellent source of hematopoietic progenitor cells. It has been widely used for bone marrow reconstitution for decades[1, 2]. The progenitor cells contained in UCB are capable of regenerating the entire lympho-hematopoietic compartment in the host. The most notable advantage of UCB transplant is the low risk of developing graft-versus-host disease (GVHD), even when donor and recipient are partially mismatched[3]. The immune cells in cord blood are virtually free from external stimulant and infection and thus were in a relatively more naïve stage. Such immunological immaturity is the key to alleviate the severity of GVHD by decreasing the alloreactive potential of lymphocytes[2, 4]. These advantages collectively expand the clinical potential of UCB transplant in many cases including some fatal diseases. The major limitation of UCB transplant, however, is the limited and inconsistent cell dose. It has been shown that the success rate of engraftment was critically dependent on the number of nucleated cells in the donor UCB[4-6].

Although UCB is now widely used for important clinical applications, we know surprisingly little about its cellular and molecular characteristics. Especially, the composition of progenitor, lymphocyte and other nucleated cells that affect the reconstitution potency after UCB engraftment is poorly understood. Recent advances in single-cell transcriptomics technology enable the exploration of cellular heterogeneity and deduction of functional relevance[7, 8]. Single-cell RNA-seq (scRNA-seq) studies of human peripheral blood (PB) cells have revealed new insights into immune cell composition and disease-related functional abnormalities[9-11]. Some works have been done about precursor cell, erythrocyte and certain T cell subtypes in single cell level, but only focus on the certain cell type or in mouse. sequencing the all cell type simultaneously may provide global information about regulation between cells[12-15]. However, scRNA-seq studies have not thoroughly characterized nucleated cells in UCB, despite their profound clinical significance. Thus, the purpose of the current study is to investigate the nucleated cells present in UCB and depict a landscape view of the cellular composition and their transcriptomes. Such key information will undoubtedly facilitate the clinical innovation to develop more efficient and cost-effective UCB transplant treatments.

## RESULT

### A single-cell transcription atlas of nucleated cells in umbilical cord blood

To acquire a cellulome map of UCB cells at single-cell resolution, we collected UCB from two donors and isolated nucleated cells for single-cell RNA-sequencing. We generated transcriptomics map of 8,947 and 10,105 single cells for the two UCB samples, sequenced to the median depth of 483,117 and 265,714 mapped reads per cell, respectively. After careful quality control and filtering by multiple criteria, single-cell data were adjusted using SVA package to minimize batch effect. As a result, we generated expression matrix robustly measuring 1,223 genes on average in each cell. Next, to identify cell populations based on their expression signatures, we computed 15 significant principle components and performed unsupervised clustering using the Seurat pipeline. Cells were visualized in two-dimension space by t-distributed stochastic neighborhood embedding (t-SNE). To determine the unique cell subpopulations and the specific state of gene expression in UCB, we utilized the public single-cell transcriptomics dataset of peripheral blood cells for comparison. This PB dataset includes 12,000 single-cell profiles of peripheral blood mononuclear cells (PBMC) measuring 1,257 genes per cell on average, which are at comparable level with those of the UCB data.

A global view was generated to illustrate the landscape of cell composition in UCB. Eleven distinct cell populations were clustered based on their gene expression profiles in both UCB samples. Public data for PB cells were processed in parallel and clustered with UCB cells in the same dimensional reduction space (Fig. 1A). To examine the inter-individual variability and investigate whether we can capture the cellular heterogeneity of all cell subpopulations in the UCB, we compared the classification results from the two donors. The identified cell clusters were quite consistent, with all the 11 clusters found in both donors. No significant residual batch bias was observed between the two UCB samples (Supplementary Fig. 1). These results demonstrate the robustness of UCB cell classification in our data. Clusters of cells that express known markers of major immune cell types were assigned with their respective identities (Fig. 1B). All 9 major immune cell types and hematopoietic lineages found in PB were identified in UCB, while two cell types, namely nucleated red blood cell (NRBC) and granulocytes, were only present in the UCB samples. While it’s well known that NRBC are universal in UCB and scarcely present in PB, the discrepancy of granulocytes are mostly due to unidentical cell enrichment experiments (Methods). The abundance of the common cell types also varied in PB versus UCB, especially in the myeloid populations, suggesting a specific immunological capacity of UCB (Fig. 1C). Furthermore, the gene expression profiles of each common cell type in UCB showed various degrees of distinction compared to those in PB. A few differentially expressed genes in B cells, NK cells, CD14^+^ monocytes and T cells illustrated the potential variations in innate immune signaling pathway activation (i.e. S100A8/9) and biological processes involves in cell proliferation, and survival (i.e. JUN/FOS pathway) at cell population level (Supplementary Fig. 1B). These differences at the population level provide a rich resource to complement transcriptomics analysis performed in bulk samples, as well as a guide to functional investigations.

**Figure 1:**
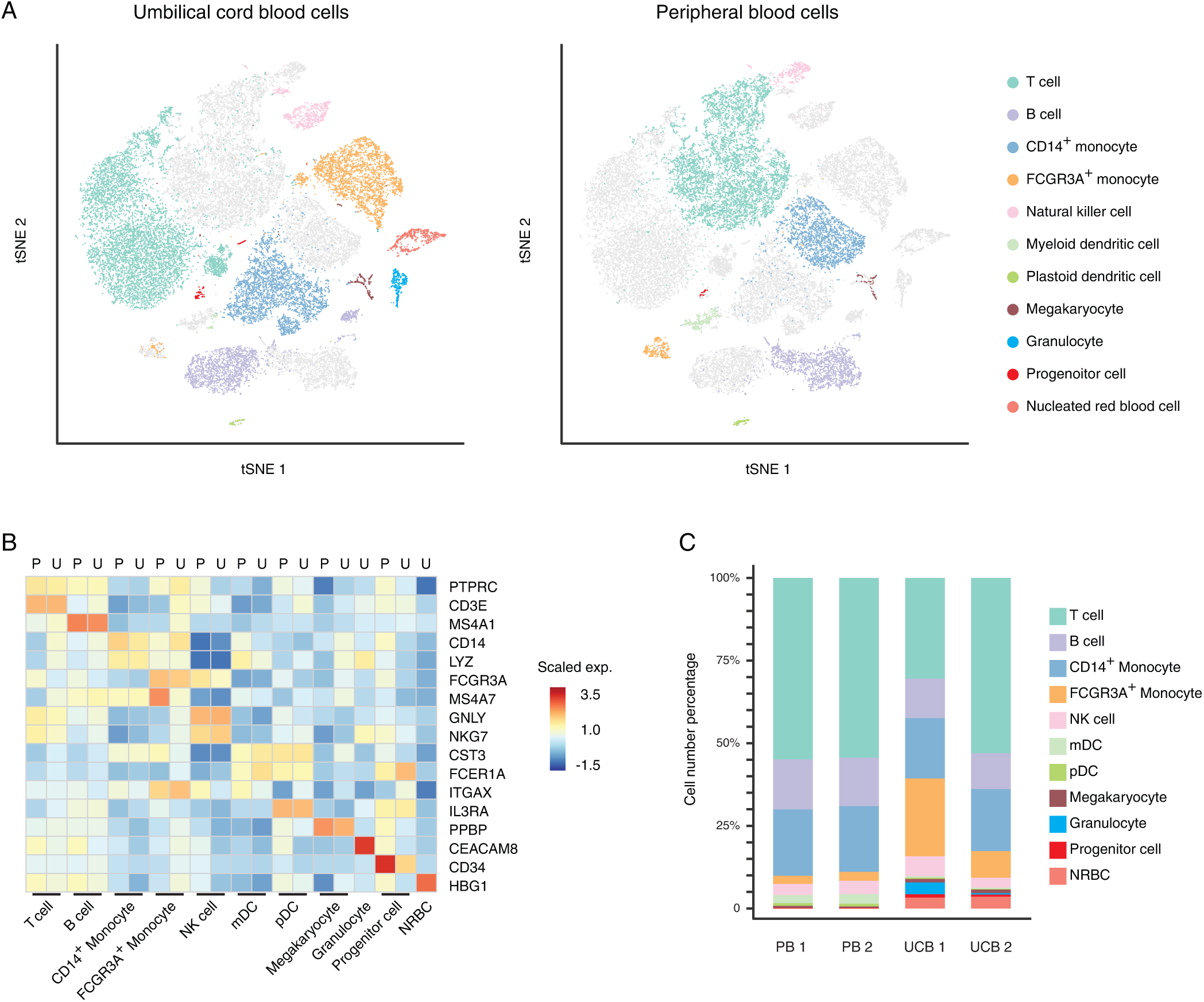
Different cell types and proportion in PB and UCB. A. Two-dimensional t-SNE distributions of UCB cells and downloaded PB cells in the same space and each cluster was assigned with their respective identities according the known markers; UCB cells were colorized on the left and PB cells were colorized on the right; All clusters were dyed at the same color according to the corresponding cell types. B. Scaled average gene expression of the major known markers (rows) detected in different cell types in UCB (U) and PB (P) (column). C. Cell type proportions of PB1, PB2, UCB1 and UCB2, which dyed using the same color in Figure 1A

### Polarity of cord nucleated red blood cell

In mammal hematopoiesis, nucleated red blood cells (NRBCs) precursors, or erythroblast, undergo several developmental stages in the bone marrow and progressively decrease cellular volume and RNA content, while accumulating specific functional proteins such as hemoglobin[16, 17]. It has been known for decades that erythroblast exist in relative large numbers in cord blood[18-20]. However, little was known about whether such development processes exist in the cord blood or whether the cord blood erythroblast population was homogenous. In our dataset, we found that NRBCs constitute a significant proportion of the total UCB nucleated cells. Interestingly, the NRBC in the UCB samples displayed pronounced polarity defined by the divergent expression of a gene repertoire. By ordering NRBCs with differential genes identified within the cluster, we deduced a pseudotime axis that suggests a gradual change of cellular state. Evidently, the NRBC formed a linear trajectory along the pseudotime axis with no significant branching, indicating that the cell polarity resulted from a continuous changes of gene expression (Fig. 2A). Next, we modeled gene expression along this trajectory to identify genes characterized by a wave-like pattern. The most prominent of these were the genes encoding surface markers and proteins critical to the function of red blood cells, such as CD47, CD36, hemoglobin and glycophorins (Fig. 2B)[21]. The CD47 molecule has long been considered as one of the cell surface markers of primitive erythrocytes[22]. Hemoglobin genes, in contrast, are highly expressed in the relatively mature form of the erythrocyte. Thus, the polarity observed most likely reflected the maturity state of the NRBCs. An intermediate cell state that bridges the naïve state (CD47 high) and the mature state (hemoglobin high) was also observed. This intermediate stage was characterized by the elevated expression of a set of genes including those encoding glycophorins, suggesting that the cells in this stage exerted a specific function, rather than just transient intermediates. Strikingly, several key transcriptional regulators of erythrocyte homeostasis, including GATA1/2 and BCL11A[23-25], also clearly exhibited divergent patterns along the pseudotime axis (Fig. 2C). GATA1 is a well-characterized transcription factor responsible for the activation of multiple hemoglobin encoding genes in erythroid ontogeny[26], while BCL11A is a transcription factor silencing hemoglobin encoding genes[25]. Another example was CITED2 and SOX6, transcription factors recently characterized as signature molecules specifically expressed in mouse primitive and definitive erythroblasts respectively, showed similar specificity in the naïve and intermediate cellular states as defined by the pseudotime axis[27]. In addition, a gradual decrease in the numbers of RNA molecules (represented by UMI) and expressed genes across the pseudotime axis was observed, supporting the correlation between linear polarity and the cord blood NRBCs functionality (Fig. 2D). These lines of evidence further corroborated the polarity identified in the NRBC population in UCB, and strongly indicated that the differential activation of transcriptional programs was one of the underlining mechanisms.

**Figure 2:**
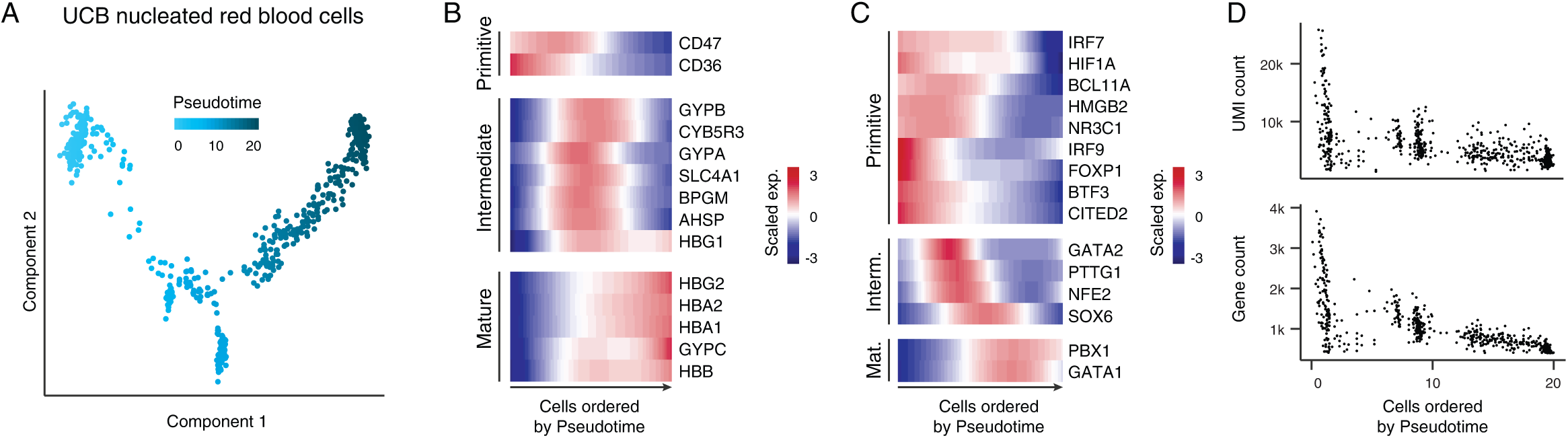
NRBCs in the UCB samples displayed a pronounced polarity. A. The ordering of NRBCs along pseudotime in a two-dimensional space determined by Monocle2. Each dot represents a single cell, and the color represents the order of pseudotime. B. Expression of genes that encoding surface markers and proteins critical to the function of red blood cells in NRBCs ordered by pseudotime of three states: Primitive stage(top), Intermediate stage(middle), Mature stage(bottom). C. Expression of transcriptional regulators in NRBCs ordered by pseudotime time of three states: Primitive stage(top), Intermediate stage(middle), Mature stage(bottom). D. Detected UMI numbers and Gene numbers in each cell ordered by pseudotime, each dot represents a cell.

### Molecular signatures of UCB progenitor cell

A distinct progenitor population was found in the UCB that shared a similar transcriptome profile with the hematopoietic stem cells (HSCs) in the PB dataset. However, when the t-SNE clustering was performed with the progenitor population in a finer resolution, a secondary subpopulation emerged, demonstrating the heterogeneity of progenitor population in the UCB (Fig. 3A). One subpopulation of UCB progenitor cells overlapped with HSCs in PB and specifically expressed the canonical HSC marker genes such as CD34, SOX4 and CD135 (red dots, Fig. 3A), suggesting their identity as being cord blood HSCs. Interestingly, the other subpopulation consists cells only from the UCB (blue dots, Fig. 3A) and did not express the HSC canonical markers (Fig. 3B, 3C) despite the similarity in overall gene expression spectrum, which drove the t-SNE clustering. Surprisingly, this CD34^-^ UCB specific progenitor population highly expressed the myeloid lineage-specific gene MS4A3 (Fig. 3C), a known signature of granulocytic-monocytic progenitors (GMPs)[28]. GMPs give rise to mast cell progenitors (MCP) and basophil progenitors (BPC), which are found in the bone marrow, spleen and gastrointestinal mucosa[29]. Furthermore, FCER1A, the gene encoding the Fc fragment of the IgE receptor, which is a surface marker frequently used in cell sorting for mast cells[30], was highly expressed in the CD34^-^ cell population; while CCR3, a sorting marker for basophils[31, 32], was co-expressed at a comparable level. Similarly, many genes that play regulatory roles in mast cell and basophil differentiation, exemplified by ITBG7 and CSF2RB[29, 33], were co-expressed at high level as well (Fig. 3C). The collective activation of gene repertoires critical in GMP-MCP and GMP-BPC ontogeny axes strongly suggested that these cells were bi-potent progenitors or intermediate cells, similar to the basophil/mast cell progenitor (BMCP) first verified in spleens of adult C57BL/6 mice[34]. High expression levels of GATA2 and median expression of CEBPA transcription factors were also consistent with the signatures of mouse BMCP [34-36] (Fig. 3C). A critical difference between the UCB subpopulation and the mouse BMCP is that CD34 expression was turned off, suggesting limited stemness in these cells. We thus hypothesized that these cells represent the intermediates before the bifurcation during basophil and mast cell differentiation and termed them umbilical intermediate bi-potent cells (uIBC).

**Figure 3:**
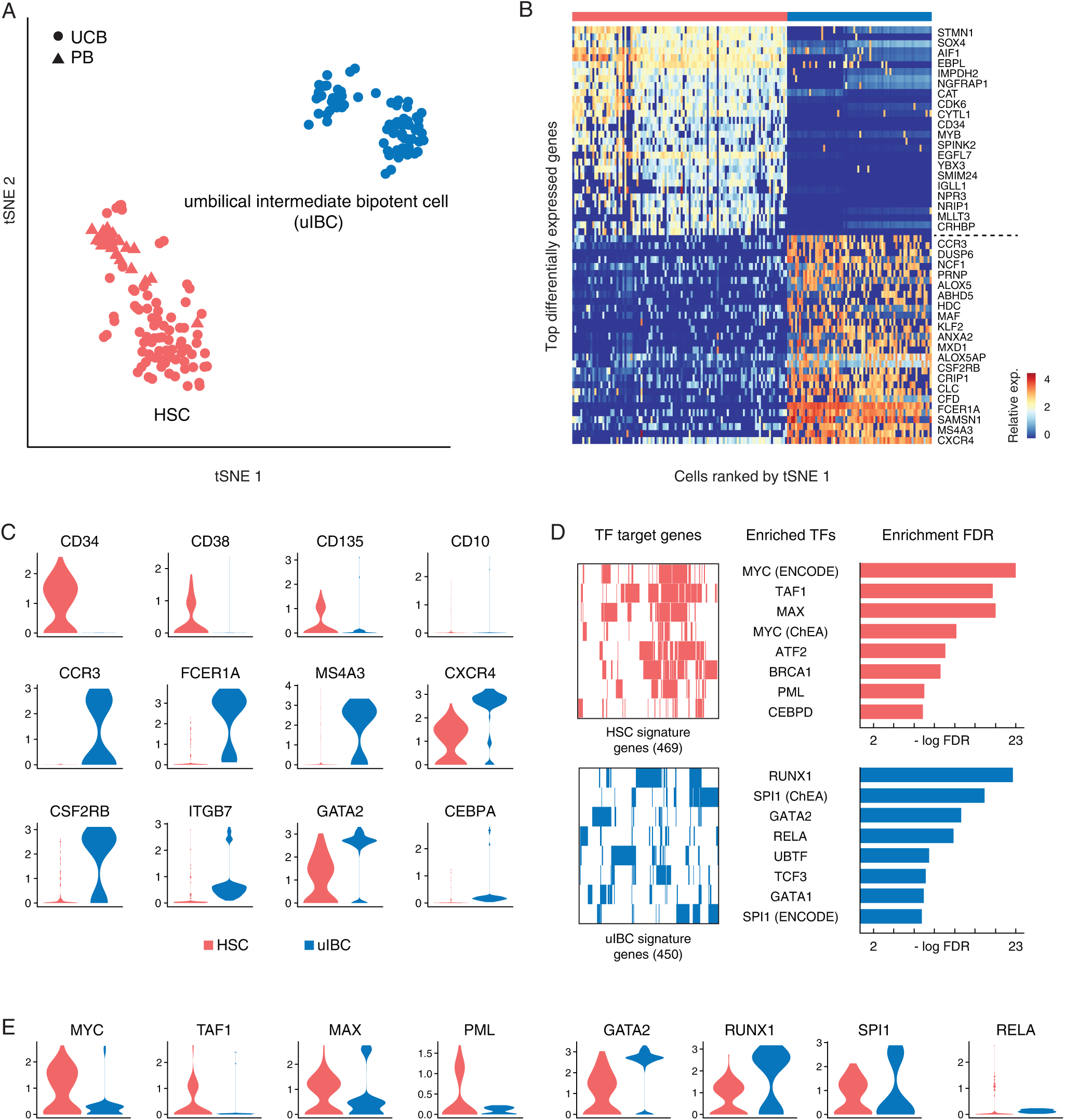
Progenitor cells in UCB samples displayed heterogeneous molecular signatures. A. The zoomed t-SNE projection of progenitor cells from UCB and PB samples, showing the formation of two main clusters shown in different colors, blue: uIBC and red: HSC. The functional description of each cluster is determined by the gene expression characteristics of each cluster. B. Heatmap of progenitor cell clusters with the differentially expressed signature genes. Cells along lateral axis were ranked by t-SNE1 in figure 3A. The panel above the heatmap identifies the different clusters. C. Violin plots show the single-cell gene expression pattern of two clusters, blue: uIBC and red: HSC. D. Transcription factor enrichment analysis of the two clusters using the HSC signature genes (469, top left) and the uIBC signature genes (450, bottom left) revealed enriched transcription factors of HSC (top middle) and uIBC (bottom middle) and corresponding enrichment score (-log(FDR), top right and bottom right). E. Violin plots show the single-cell transcription factors pattern of two clusters, blue: uIBC and red: HSC.

A spectrum of differentially expressed genes was identified in the HSC and uIBC subpopulations in UCB (partly shown in Fig. 3B). Next, we asked whether the switch of cell identities resulted from alteration of transcriptional programing that governed the differentiation process. Transcription factor enrichment analysis utilizing the comprehensive database of Encode[37] and ChEA[38] was used to detect over-represented combinations of conserved transcription factor binding sites in a given set of genes. The analysis revealed that MYC, TAF1 and MAX were the mostly enriched for activating highly expressed genes found in the HSCs as compared to uIBC. These transcription factors are well known for their roles in proliferation and cell cycle control[39-41]. Conversely, RUNX1, SPI1 and GATA2 were ranked as the top enriched transcription factors for activating the highly expressed genes in the uIBCs (Fig. 3D). These transcription factors are conventionally considered as master regulators of differentiation of the myeloid lineage[35, 42, 43]. Such functional correlation was further corroborated by the mutually exclusive expression pattern of the top enriched factors. For example, high expression levels of MYC, TAF, MAX and PML, enriched for activating HSC feature genes, were detected in the HSCs; and *vice versa,* high expression levels of GATA2, RUNX1, SP1 and RELA, were detected in the uIBC (Fig. 3E). These data collectively supported the notion that the two subtypes of cells we found in the progenitor population in UCB were indeed functionally relevant.

### Heterogeneity of natural killer T cell in UCB

Effective immune response against infection, allergy and cancer generally requires coordinated activation of both the innate and adaptive immune systems. Recent studies have shown that natural killer T (NKT) cells emerge as a bridge between innate and adaptive immunity in order to mediate an immune response[44]. Unlike conventional T cells, NKT cells exhibits distinct tissue specificity under homeostatic conditions, suggesting compartmentalized functions[45-48]. Given the immune immaturity of cord blood cells, we expected an unique pattern of the NKT population. We selected all NK cells and the T cells that express *NKG7* as well as the T cells nearby on the t-SNE space in Fig. 1A from PB and UCB datasets and performed further t-SNE clustering to systemically examine their potential heterogeneity. As a result, these cells clustered into five distinct subgroups in PB and six subgroups in UCB (Fig. 4A). By analyzing the expression of the lineage marker genes *CD3* and *KLRB1,* we assigned the cell clusters with T (CD3^+^KLRB1^-^), NKT (CD3^+^KLRB1^+^) and NK (CD3^-^KLRB1^+^) lineages[46, 49, 50] (Fig. 4A). Interestingly, apparent heterogeneity was observed in both T and NK cells. For example, the CD3^+^KLRB1^-^ T cells in both PB and UCB could be further sub-grouped into cytotoxic and non-cytotoxic populations, based on the expression of cytotoxic marker genes, such as *GNLY, GZMK, GZMB* and *PRF1.* Similarly, NK cells could also be sub-grouped into GZMK^+^ and GZMB^+^ populations, based on the mutually exclusive expression pattern of the two granzyme genes. However, no significant difference in gene expression was detected between UCB and PB NK cells, indicating that the GZMK^+^ and GZMB^+^ NK populations were both constitutive of general homeostasis.

**Figure 4:**
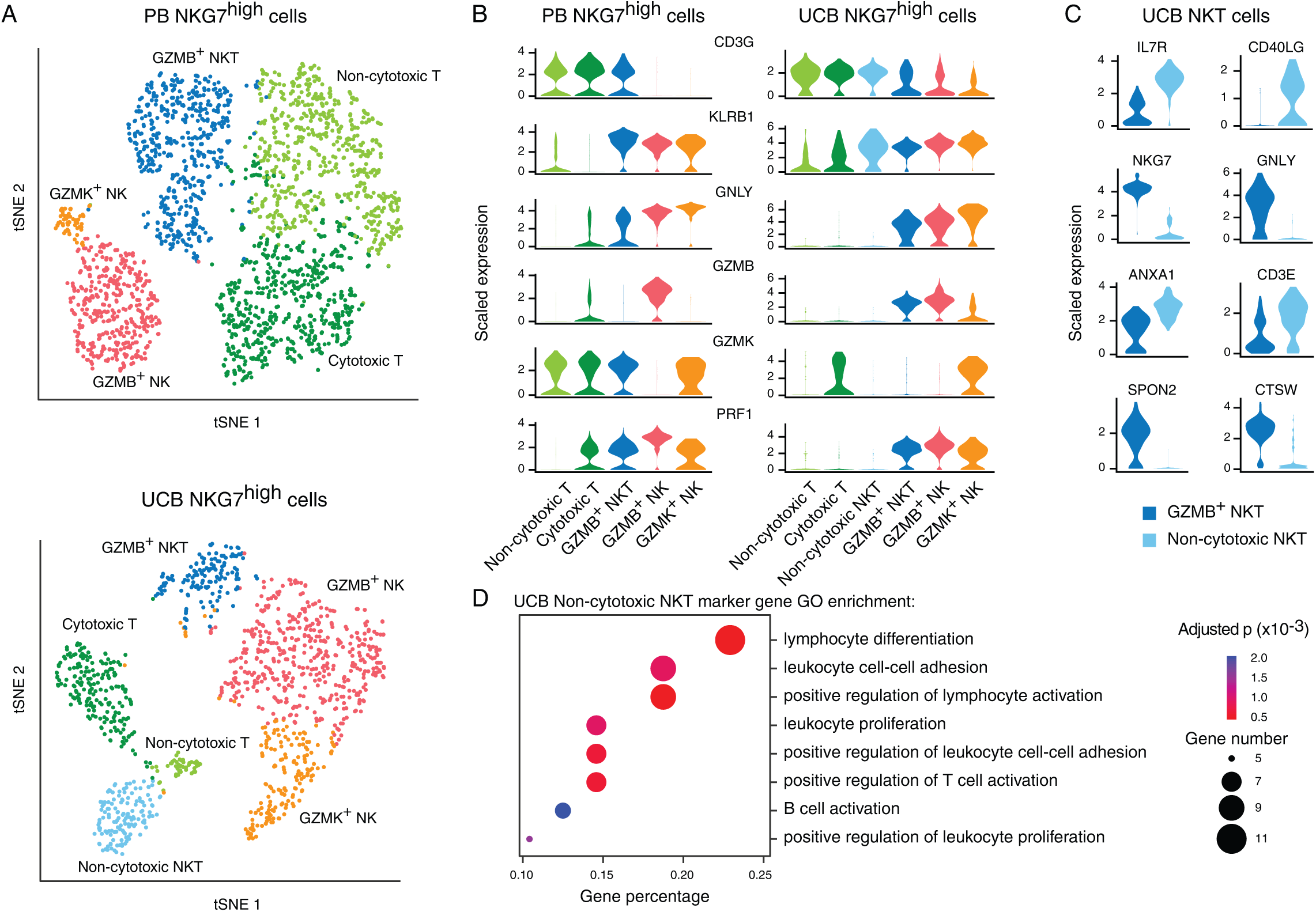
NKT cells and NK cells showed functional heterogeneity in UCB and PB. A. Two-dimensional t-SNE distributions of all NK cells and the T cells that express NKG7 gene from PB and UCB datasets. Color represents five distinct subgroups (GZMB^+^ NKT, GZMK^+^ NK, GZMB^+^ NK, Non-cytotoxic T, Cytotoxic T) in PB (top) is and six subgroups (GZMB^+^ NKT, GZMK^+^ NK, GZMB^+^ NK, Non-cytotoxic T, Cytotoxic T, Non-cytotoxic NKT) in UCB (bottom) according corresponding cytotoxic signature genes. B. Violin plots show the single-cell gene expression pattern of signature genes between the distinct subgroups in UCB (right) and PB (left). The shapes represent the distribution of cells based on the scaled expression. Colors represent the different subsets. C. Violin plots show the scaled single-cell expression of indicated differentially expressed genes between GZMB^+^ NKT and Non-cytotoxic NKT subsets in UCB. D. Gene ontology analysis of GZMB^+^ NKT and Non-cytotoxic NKT subsets in UCB using the differentially expressed genes. X-axis represents the enriched gene percentage in certain pathways. Y-axis shows the enriched pathways ordered by gene percentage. The size of dot represents the gene numbers. Color scale shows the adjusted p-values of each enriched pathway.

NKT cells, originally clustered as parts of the T cell population in PB and UCB, express high levels of GZMB and PRF1 but not GZMK. Interestingly, we found a unique subpopulation of NKT cells (non-cytotoxic, light blue dots in Fig. 4B) in UCB that do not express the GNLY, GZMK, GZMB or PRF1 genes, suggesting the lack of cytotoxicity mediated by the granzyme and perforin pathway (Fig. 4B). NKT cells were previously reported to have tissue-specific gene expression programs that lead to diverse functions and were termed NKT1, NKT2 and NKT17, predominantly localized in liver, lung and peripheral lymph node, respectively[46, 51-54]. In our data, the expression profile of the GZMB+ NKT cell was mostly similar to that of the NKT1 type, highlighted by signature expression of IL2RB, KLRB1, ZBTB16 and TBX21 (Supplementary Fig. 2A), but these cells did not express GATA3[55]. The non-cytotoxic NKT cell was not a part of NKT2 and NKT17, because they exhibited the expression pattern of high level of KLRB1 and GATA3[55, 56], indicating a new subtype of NKT cell in UCB. We then considered whether the development stage that shapes the different clusters, but all these two NKT cells are belong to CD24^-^CD44^+^KLRB1^+^ stage 3 (Fig. 2B, Supplementary Fig. 2A). In contrast, the non-cytotoxic NKT cells highly express a spectrum of surface molecules, such as IL7R, CD40LG and CD3E that may mediate interactions with other immune cell types to coordinate immune response (Fig. 4C). Gene ontology analysis further corroborated that the highly expressed genes of the non-cytotoxic NKT cells were enriched in lymphocyte cell-cell adhesion pathways and lymphocyte proliferation, differentiation and activation processes (Fig. 4D). Thus, we conclude that the cell composition of NKT and other cytotoxic cells vary between PB and UCB.

## DISCUSSION

For the first time, we present here a single-cell level transcriptomic landscape of nucleated cells in UCB. By analyzing the expression pattern of known marker genes, we identified UCB cells belonging to almost all of the major hematopoietic lineages in the PB, covering lymphoid, myeloid and hematopoietic progenitor cells. We also observed that certain cell populations were highly enriched in UCB cells, such as the NRBCs and the uIBCs. In adults, red blood cells are generated mainly in the bone marrow from nucleated cells identified as erythroid precursors. These cells undergo morphological changes through cell divisions and gradual decrease in cell size and RNA species, increase in chromatin condensation and hemoglobin protein accumulation. Such changes have been associated with the early stages of maturation of red blood cell. Here, in our dataset we also observed such a dynamic cellular state in a linear polarity, suggesting that erythroid precursors undergo a similar maturation process in the UCB as in the bone marrow. Progenitor cell populations in UCB also appeared to be a mixture of two distinct subpopulations. The uIBC, a unique subpopulation only seen in UCB but not PB, were identified with characteristics of both basophil and mast cell signatures. A similar bipotent population (BMCP) exists in mouse spleen and is capable of divergent development. Signature gene expression, including transcription factors and surface markers were remarkably similar between BMCP and uIBC, except that uIBC lack the expression of the conventional progenitor marker CD34. Although uIBC and HSC in UCB were globally similar in their transcriptomic profiles, the lack of CD34 made it difficult to conclude whether these uIBCs were indeed progenitors or transient intermediates captured during UCB hematopoiesis. Functional validations will be necessary to determine the potential abilities of self-renewal and lineage regeneration of these cells and substantiate the similarity with mouse BMCP at the functional level. Next, we discovered that gene expression signatures often vary greatly in the common cell types seen in both UCB and PB. We interrogated the UCB single-cell data at a finer scale and discovered a previously unknown NKT population that may be unique to UCB. NKT cells have an essential role in bridging innate and adaptive immunity against infectious diseases and tumorigenesis. Thus, NKT cells have significant therapeutic values. UCB transplants have demonstrated remarkable effectiveness in treating many types of blood cancers. Adoptive transfer of the NKT cells has been tested in animal models[57, 58], and several clinical trials are in process to test the safety and efficiency of NKT cell transfer to harness the solid tumors in human[59-62]. The enhanced understanding of the NKT cell heterogeneity in UCB would benefit our selection of appropriate source and the activation of the cytotoxicity of NKT cells to target cancer and other diseases. Therefore, we speculated that a targeted enrichment, modulation or engineering of the existing NKT populations in the UCB could lead to considerable improvement in the efficacy of enhancing protective immune responses.

Taken together, our data provides the first single-cell transcriptomic references for UCB, which could be used as a standard dataset for comparative analysis. We expect that this dataset will prove useful in uncovering the novel molecular signatures that define the cellular heterogeneity in UCB and provide markers for targeted enrichment of certain cell types of interest to researchers in multiple fields. Our dataset should also be a rich resource to formulate hypothesis of signaling pathway activation, transcription control and other mechanistic studies in the field of functional immunology in single cell level Moreover, to our knowledge, the current study was the first to performed 10X Genomics single-cell transcriptome profiling in tandem with the rolling circle amplification-based BGISEQ-500 high throughput sequencing platform, broadening the technological diversity for single-cell RNA-seq applications.

## METHODS

### Sample collection

Two umbilical cord blood samples were collected from healthy donors immediately after caesarean section with informed consents. Samples were stored in EDTA anticoagulant tubes and transported to laboratory within 1 hour. CD45^+^ and CD45^-^ cells were isolated from 1 mL cord blood by positive and negative selection, respectively, using Whole Blood CD45 MicroBeads (Miltenyi,130-090-872) and Whole Blood Column Kit (Miltenyi, 130-093-545). Next, the CD45+ and CD45- cells were counted by hemocytometer and mixed at the ratio of 4 to 1. The cells were further gently pipetted into a single-cell suspension and diluted to 700 cell/μL concentration. The single cell gene expression dataset of peripheral blood from healthy donors were downloaded from the https://support.10xgenomics.com/single-cell-gene-expression/datasets (processed by Cell Ranger 2.0.1). Peripheral blood mononucleated cells were isolated and processed in 10X platform (10X Genomics, USA).

### Library construction and sequencing

Single-cell suspension was loaded to Single-cell 3’Chips (10X Genomics, USA) and subjected to GemCode Single-Cell Instrument (10X Genomics, USA) to generate single-cell Gel Beads in Emulsion (GEMs), per manufacture’s instruction. GEMs were next subjected to library construction by Chromium™ Single-cell 3’ Reagent Kits v2 (10X Genomics, USA), steps of which included RT incubation, cDNA amplification, fragmentation, end repair, A-tailing, adaptor ligation, and sample index PCR. However, such library was originally designed to be sequenced by the Illumina sequencing platform. In order to convert the libraries to that compatible with BGI-Seq 500 sequencer, we performed a 12 cycle PCR on the libraries with BGI-Seq adaptor primers, and subsequent DNA circularization, rolling-cycle amplification (RCA) to generate DNA Nano Balls (DNBs). The purified DNBs were sequenced by BGISEQ-500 sequencer, generating reads containing 16 bp of 10X™ Barcodes, 10 bp of unique molecular indices (UMI) and 100 bp of 3’ cDNA sequences. Each library was sequenced in three lanes, yielding ~1.9 billion reads in total.

### Alignment and initial processing of sequencing data

CellRanger toolkit (10X Genomics, USA, version 2.0.0) was employed to align the cDNA reads to GRCh38 transcriptome. Filtered UMI expression matrices of both samples were generated with the default parameter settings except “--force-cells=4000”[63].

### UMI normalization and cell clustering

The expression matrices of both samples were normalized by “cellranger aggr” function from the CellRanger toolkit, with the parameter “--normalize=mapped”. In order to remove potential technical bias introduced during library construction and sequencing, SVA package was used to minimize such batch effects. The Seurat package was then utilized to quality-control expression data for each cell, first by removing the undetected gene (i.e. expression in less than 30 cell), secondly by removing the cells with less than 400 detected genes. Cells that yielded high level (>10%) of reads that aligned to mitochondria genes were also removed. Next, the filtered expression matrices were used for unsupervised cell-clustering by the Seurat package, adapting the typical pipeline that was recommended by the authors[64]. Subsequently, the identity of each cell cluster was annotated manually, based on the previously known markers and the cluster-specific genes listed by the Seurat “FindAllMarkers” function.

### Pseudotime analysis

To infer the developmental polarity of nucleated red blood cells (NRBCs), pseudotime analysis was performed by the Monocle2 package (version 2.6.4). NRBCs were ordered according to the pseudotime deduced by the top 1,000 differentially expressed genes within the NRBCs.

## AUTHOR CONTRIBUTION

X.LIU. and Y.H. jointly supervised research. Y.Z. and G.Y. designed the experiments. X.W., K.G., Y.Z. and X.Z. performed the experiments. Y.Z. and J.W. processed single-cell RNA sequencing data. Y.Z., X.LI., J.W. and Z.W. analyzed the data. W.Z. and B.F. collected the cord blood. X.LI. and Y.Z. wrote the manuscript. X.LIU revised the manuscript. All authors have reviewed and approved the final manuscript.

## ACKNOWLEDGMENTS

We thank the two donors who generously provided the UCB samples. We also thank Liqin Xu, Zhikun Zhao for helpful discussions and BGI colleagues who have helped producing the high quality data. This work was supported by Shenzhen Municipal Government of China (JCYJ20170817145404433 and JCYJ20170817145428361)

---

**Supplementary Figure 1:**
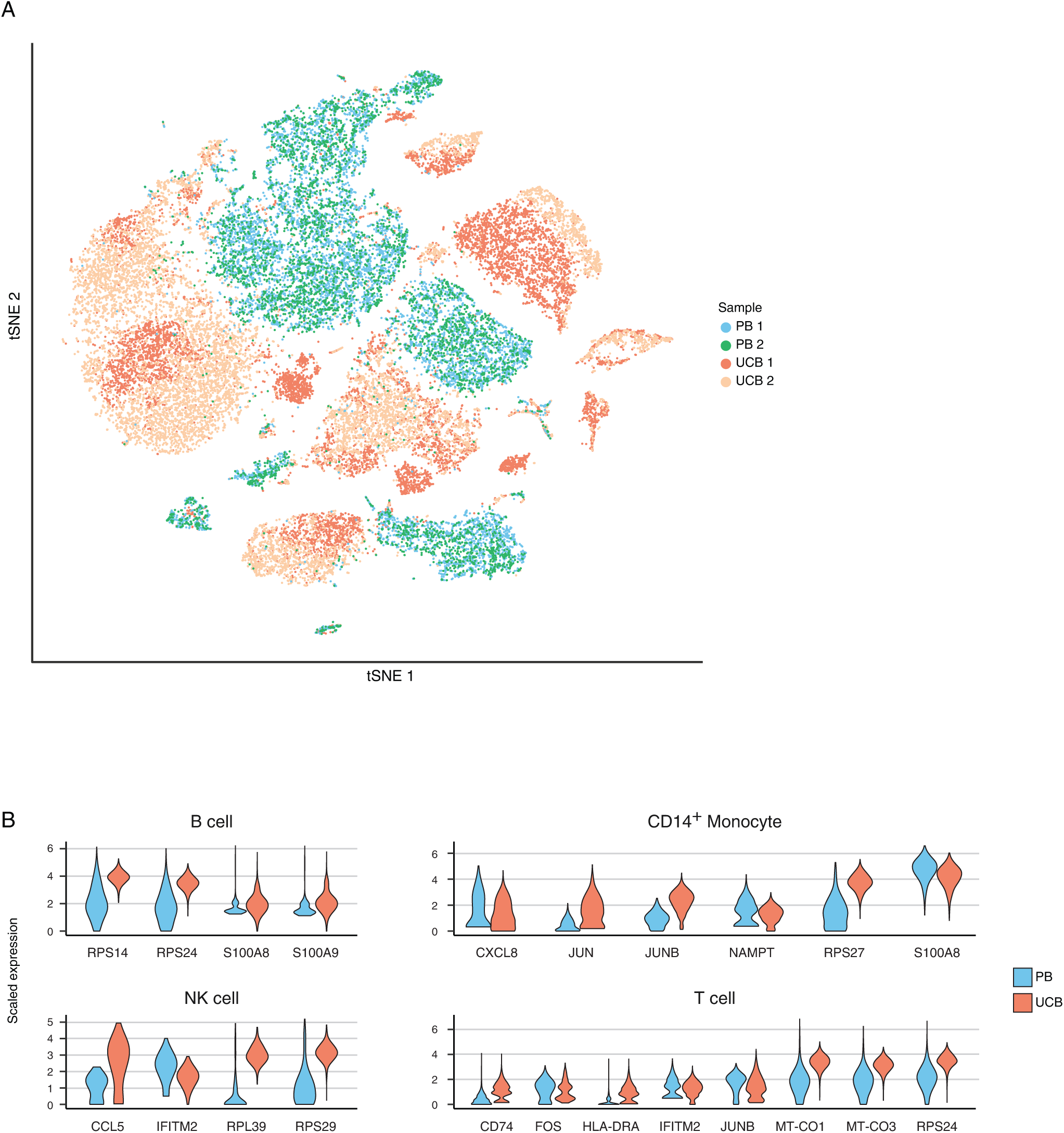
Sample distribution and gene expression differentiation between UCB and PB. A. Sample distribution in the same two-dimensional t-SNE space as Figure 1A. Each dot represents a single cell. Color represents the different samples in UCB and PB. B. Violin plots show the scaled single-cell expression of indicated differentially expressed genes between UCB and PB in B cell, CD14^+^ Monocyte, NK cell and T cell.

**Supplementary Figure 2:**
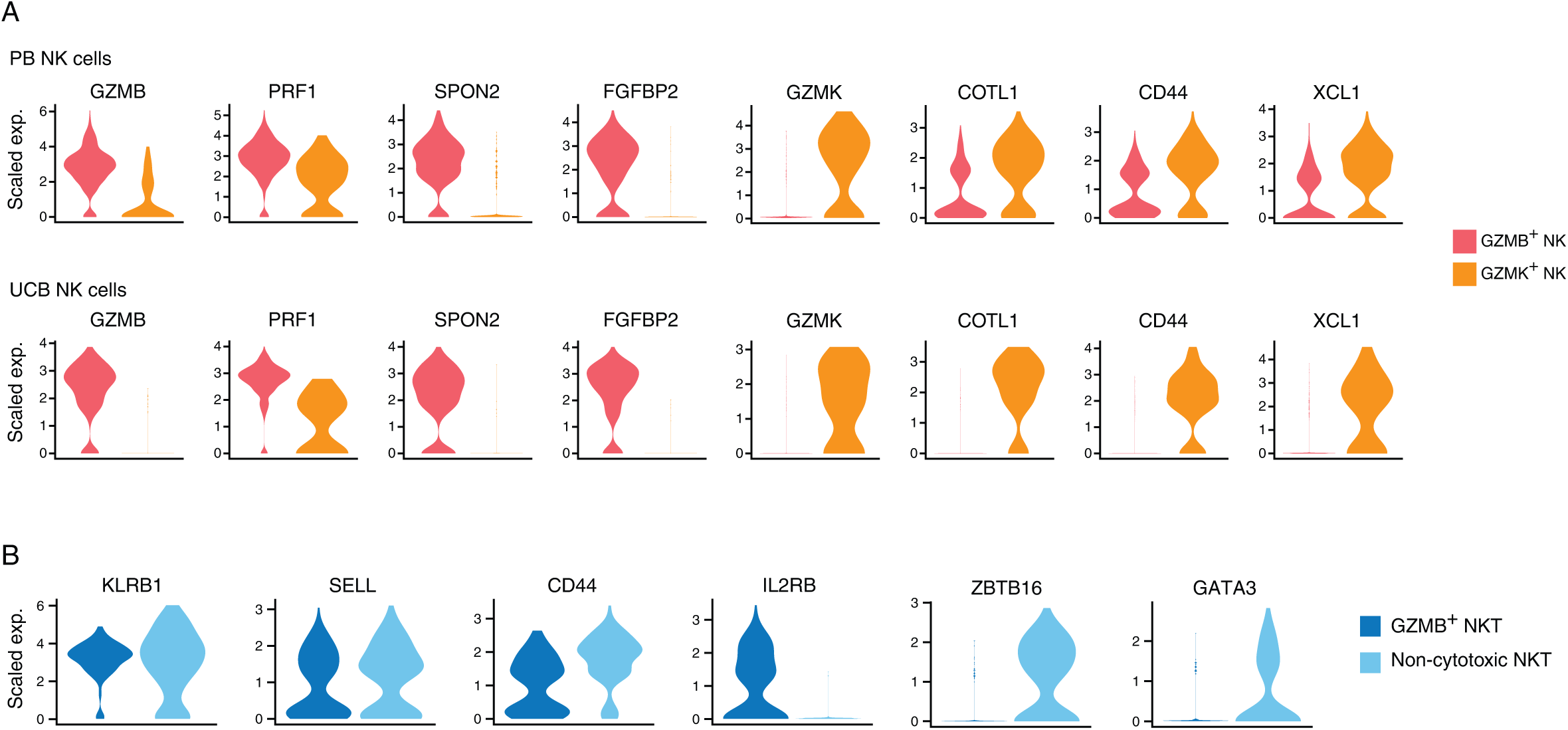
Cell subsets express distinct set of transcripts in NK and NKT. A. Violin plots show the scaled single-cell expression of indicated differentially expressed genes between GZMB^+^ NK and GZMK^+^ NK in PB (top) and UCB (bottom). B. Violin plots show the scaled single-cell expression of indicated differentially expressed genes between GZMB^+^ NKT and Non-cytotoxic NKT subsets in UCB.

